# GLORB: Robust Bayesian inference for differential expression under global expression shifts

**DOI:** 10.64898/2026.08.28.747928

**Authors:** Rowan Callahan, Stephen D. Coleman, Thuy T. M. Ngo

## Abstract

Estimating differential gene expression is a common task in RNA-seq. Current methods mostly rely on normalization to reduce variance and increase accuracy. These methods are widely used and provide invaluable information about transcriptomic changes between biological conditions. However, widely used normalization methods are known to distort estimates of transcript differential expression when a majority of genes are upregulated or downregulated, or when the total RNA content per cell changes. Despite the presence of global expression shifts in a number of contexts, few methods exist that can provide accurate normalization and estimate linear models under this context without spike-in controls. Here, we present **GLORB** (**GLO**bal-shift **R**obust **B**ayesian model), a method for estimating generalized linear models under global upregulation. We develop two models that are able to recover differentially expressed genes and linear model coefficients with lower distortion of results. We show that our model’s method of accounting for library size variance is consistent with DESeq2’s median of ratios, and edgeR’s trimmed mean of M-values under conditions when a minority of genes are upregulated and outperforms them under circumstances when most genes are either increased or decreased between groups. Finally, because our method does not rely on calculating geometric means for each gene it is able to work in datasets with much higher sparsity.

## 1 Introduction

RNA-seq has become a fundamental assay in cell biology and the most common measure of transcriptomic abundance in modern cell biology [37, 43]. RNA-seq is a developed and highly utilized methodology which is now used for drug repurposing studies [20, 39], as a readout of tissue state [12], pathway analysis [38] as well as general perturbation analysis [8]. In cancer research alone, it has been used to create tumor subgroupings [15], predict patient response to treatment, and predict disease status and prognosis [15].

A typical bulk RNA-seq experiment involves collecting tissue samples from at least two groups; typically a case and a control group, with the intention of identifying genes that are increased or decreased when the case compared is to the control. Mathematically similar data is also collected by a number of other assays that produce count based readouts such as ATAC-seq or ChIP-seq – in these cases the object of inference is differentially accessible regions or DNA regions differentially bound by proteins rather than genes. In all cases we are comparing tables of counts across multiple samples from multiple groups. Count data are characterized by variance that depends on the mean (heteroskedasticity), i.e. as the total number of counts increases so too does the variance, which demands appropriate modeling choices to reflect this. Beyond this intrinsic property, observed counts from these assays are assumed to be confounded by sources of *technical variation*; noise that come from sample handling and processing and is unrelated to the biology of interest. One of the most important of these assumed effects is called the *library size* and relates to the total amount of RNA that is analyzed once the data is sequenced. The dominant framework for addressing this confounder is to divide all observed counts within a sample by a sample-specific *normalization factor*, *s_n_* called the size factor, intended to account for processing effects that would scale up or down all counts and make samples directly comparable by removing the assumed multiplicative technical offset. This scaling model assumes that the observed counts for gene *g* in sample *n* is of the form *y_ng_* = *s_n_µ_n,g_*, where *µ_n,g_* is the underlying mean number of counts per biological unit (grams of tissue, number of cells etc.) within the sample *n* for this gene *g* which is then compared for all samples across conditions using some statistical test such as the Wald test [25].

Two methods dominate current practice in accounting for this technical variation in RNA-seq datasets: DESeq2 [25], which uses the Median-of-Ratios (MoR) estimator for *s_n_*, and edgeR [35], which uses a trimmed mean of ratios which they call M-values (TMM). These methods have also continued to be used in the era of single-cell RNA for calculations in so called pseudo-bulk analyses and are often used for both ATAC and ChIP-seq data when analyzing bulk datasets. Both estimators of the size factor are unbiased only under the assumptions that a majority of features are unchanged between conditions; that there is no systematic shift in expression or accessibility that would shift the majority of genes or the median in one condition relative to the other. Specifically, MoR assumes that the median log-ratio of counts is 0 (i.e., the median gene is not differentially expressed), and TMM explicitly trims the distribution of per-gene fold changes before computing a mean, relying on these trimmed quantities being centered at zero. When these assumptions hold, these provide an unbiased estimate of the size-factor *s_n_* for scaling the library size.

However, the validity of this assumption has been questioned since the early days of RNA-seq analysis, [26]. In such conditions where there is a global shift in regulation, then these normalization methods and common competitors are mis-specified, producing biased estimates and leading to a higher percentage of false positive estimates of increased or decreased (differentially expressed) genes [9]. Empirical evidence that this assumption is violated in many biological systems is now substantial. Lin et al. [22] showed early on that c-*MYC* amplification drives global expression changes. Subsequent large-scale analyses of the Cancer Genome Atlas (TCGA) data by Zatzman et al. [46] and Cao et al. [2] observed hypertranscription of RNA transcripts produced in specific cancers compared to controls when normalizing using a mutation based method. Global upregulation has also been reported under various plant growth conditions [1, 21] and is assumed in other conditions such as stem cell differentiation [32]. The downstream consequences of these biased estimates is not limited to incorrect fold-change estimates on a per gene basis. RNA-seq analyses are often used to prioritize pathways for further research, thus the long term effects of this offset in calculated changes could have serious knock-on effects for pathway analysis, drug target prioritization, understanding of drug effects, understanding cell state, cell typing, cancer and other disease sub-typing, and understanding tissue state. A systematic bias in the direction of differential expression has the potential to propagate through all downstream analyses and decision-making.

Various computational and biological methods exist that partially address the problem of global upregulation. Spike-in controls are synthetic RNA molecules of known concentration that, when correctly added to each sample before library preparation, provide an external reference to normalize technical variation against [1, 17]. A major validated source of these come from the External RNA Controls Consortium and are often referred to as ERCC spike-ins. These are considered the gold standard for removing library-size effects from within samples and can address global upregulation; however, using spike ins effectively requires extremely careful liquid handling to ensure effects between samples are kept constant and still require exact and standardized human handling between conditions. Importantly, they must also be integrated in from initial experimental design and many existing public datasets lack such controls. On the computational side, tools such as qsmooth [14] which performs quantile smoothing within conditions and ISnorm [23] which predicts stable genes when a large number of cell replicates is available. However ISnorm is designed for single cell analysis with 1000s of cells to find stable genes and qsmooth cannot be used with a full linear model and design matrix precluding the usage of variables like age or variables with interaction terms. Other methods find control genes and then extrapolate them back to previous samples that were processed without spike-in data use techniques such as quantitative PCR (qPCR) [30] or orthogonal measurements such as estimating tumor fraction from DNA mutations in order to calculate global changes in RNA expression [2, 46]. All these methods can be used to find conditions that have global upregulation where a majority of genes are increased between samples. However, to our knowledge there is no computational tool that exists that can process RNA-seq or other counts data and provide the same benefit of spike-in controls such as ERCC spike-ins while providing the same flexibility of linear models to any sequenced bulk RNA dataset.

To this end we attempt to bridge this gap with a Bayesian generalized linear model (GLM). We turn to the Bayesian paradigm as such models are well-established in their application to complex biomedical data, e.g., [5, 7, 13, 19], and allows us to describe our prior beliefs about the relationship between biological and technical variation through a hierarchy of latent variables [4, used such a framework in the context of ELISA data]. In this manuscript we introduce our models to address the problem of biased inference in systems with global upregulation of gene expression or accessibility, **GLO**bal-shift **R**obust **B**ayesian model, **GLORB**. Our models provide uncertainty estimates relating gene expression changes to any linear matrix of covariates, while also remaining stable under global increases in expression and gene regulation when compared with current methodologies in the field.

Our models enable identification of differentially expressed genes even within the global upregulation context provided that sample condition group assignment is not strongly correlated with batch or processing group assignment; i.e., that samples from distinct biological groups are processed in an interleaved or randomized fashion. If processing and library effects are expected to be exchangeable then our non-size-factor model is the suggested model for users as it makes this assumption but has potentially higher performance in extreme conditions. If this condition is expected to hold more weakly (for example treatment and control are processed separately but in a similar manner), we make available a second model we call our “size-factor” model. This model is designed to only require that a significant portion of genes (more than 20%) are not changed across condition. This enables principled analysis in the large body of public RNA-seq data for which spike-in measurements are unavailable.

In the remainder of this manuscript, we describe the models, and show our simulation study and empirical results from an ERCC spike-in dataset and existing TCGA data. We include a single ATAC-seq analysis to highlight the generalizability of the method, while focusing on RNA-seq generally.

## 2 The GLORB Model

We present a new method for inferring differential expression when global expression changes exist we call **GLO**bal-shift **R**obust **B**ayesian modeling or GLORB. We propose two sub-models. The first, and most similar to previous approaches, we refer to as our *size-factor model*, assumes that when comparing groups or conditions there exists a subset of genes that are not affected by model covariates. This is a much weaker assumption than the standard assumptions of TMM, center log ratio (CLR), and MoR normalization strategies–we require that a subset of genes are unaffected by condition/covariates rather than requiring a majority. However, to address the extreme case where these assumptions are violated and every gene is affected between groups/conditions, we propose our *non size-factors model* that assumes that the size-factors for each sample are fully independent of condition or group within the design matrix.

Both of these models have a sparsity inducing prior placed on them in the form of either a horseshoe prior [3, 34] in our non-size-factor model or a mixture model that is similar to a spike and slab prior [10, 28] in our size-factor model. This is important for regression models in situations where the number of covariates *P* is much larger than the number of samples *N* (*P ≫ N*) and prevents overfitting the model.

### 2.1 Model overview

The basis for our models is the generalized linear model; specifically using a Negative Binomial likelihood and a log link function as is typically used when modeling over-dispersed count data and is used by edgeR and DESeq2. The log mean *η* is described by

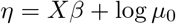

where *X*, is the design matrix of dimension *N × F* with entries *x_n,f_* for each sample *n* and factor *f*; *β* is the coefficient matrix of dimension *F × G* with entries for every factor and gene *β_f,g_*; and *µ*_0_ is the base mean–this parameter could be absorbed into *X* but we keep it separate as this coefficient is used for downstream calculations in our non-size-factor model. The mean matrix *µ* of dimension *N × G* is

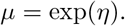

The observed data is then assumed to be sampled from a negative binomial; we use the mean-dispersion parameterization, i.e.,

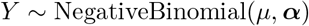

where

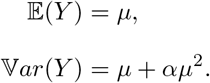

***α*** is referred to as the over-dispersion parameter and is strictly positive.

As a baseline shared by both our models, we assume that every gene has a unique over-dispersion parameter.

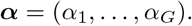

Given this context, we can now describe the key conceptual differences between our two models. Each model explores a different answer to the question, should technical variance be accounted for in the mean or the over-dispersion?

In our first model, the size-factor model, we decompose the mean parameter to include per-sample library size effects as in DESeq2 and edgeR with a sample specific library-size factor, *s_n_*. As described in the introduction, this is a quantity that inflates the expected count of every gene within the *n^th^* sample in a multiplicative fashion, and the likelihood of a count *Y_n,g_* in this model becomes:

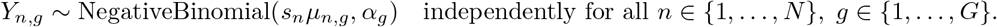

Note that while at this level our model is highly similar to these existing methods, it is differentiated as all model parameters are simultaneously inferred such that uncertainty from the size factor *s_n_* is propagated to be included in the uncertainty on the log fold changes and through our inclusion of priors that help solve the overspecification problem on coefficients and induce sparsity in our model of log fold changes.

In our second model, which we dub our non-size-factor model, we decompose the over-dispersion further into a *G*-vector and an *N* -vector; i.e., each gene has a unique entry as does each sample. To avoid notational ambiguity, we denote the per-sample over-dispersion by ***ϕ***, thus we have two vectors

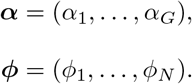

In this model we view library-size effects from the perspective of excess dispersion rather than a shift in the mean; that is, a multiplicative increase in the mean from size factor scaling should generally increase the observed variance about the true mean unless the scaling factor is exactly 1. We simplify the way that these per-gene and per-sample over-dispersion terms interact such that the over-dispersion about the mean for the *g^th^* gene within the *n^th^* sample is

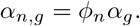

This multiplicative decomposition assumes that observed variance from library size effects and variance in counts that is inherent to each gene behave independently from one another. Thus, the count *Y_n,g_* for the *g^th^* gene in the *n^th^* samples in this model is:

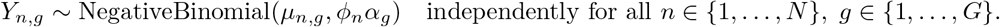

Our full hierarchical model for the size-factor model is:

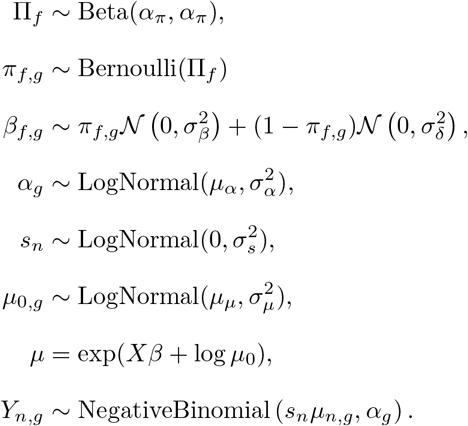

Here there is a continuous spike-and-slab prior similar to [10] on the entries of the coefficient matrix, *β*, to induce sparsity, and 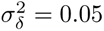 for numerical stability. The spike here represents coefficients for genes that do not change between conditions, and could potentially be considered housekeeping genes etc.

Our non-size factor model is defined

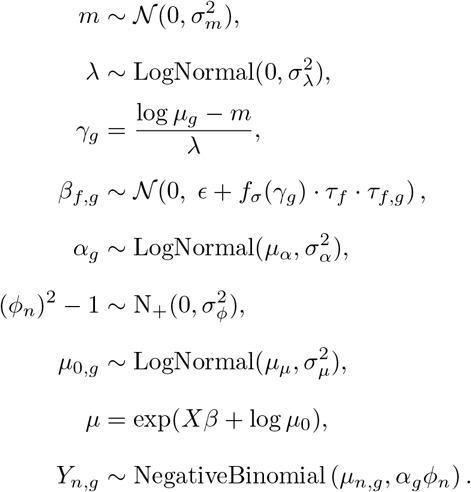

Here *N*_+_ denotes a half-Normal distribution and (*ϕ_n_*^2^ *−* 1) ensures the sample-specific over-dispersion parameter can only inflate, not deflate the over-dispersion due to *ϕ*. *f_σ_* is our notation for the sigmoid function; we include this sigmoid function for genes with low counts to avoid diverging log fold changes when counts approach zero. The entries of the coefficient matrix, *β_f,g_* have a modified regularised-horseshoe prior [3, 34] to induce sparsity. As in the previous model, genes who have zero coefficients in their fold changes can be viewed as housekeeping genes or other genes that are constant between conditions.

A graphical model depicting these hierarchies is shown in Figure 1, and the choices of hyperparameters are shown in Tables 1 and 2.

**Figure 1.**
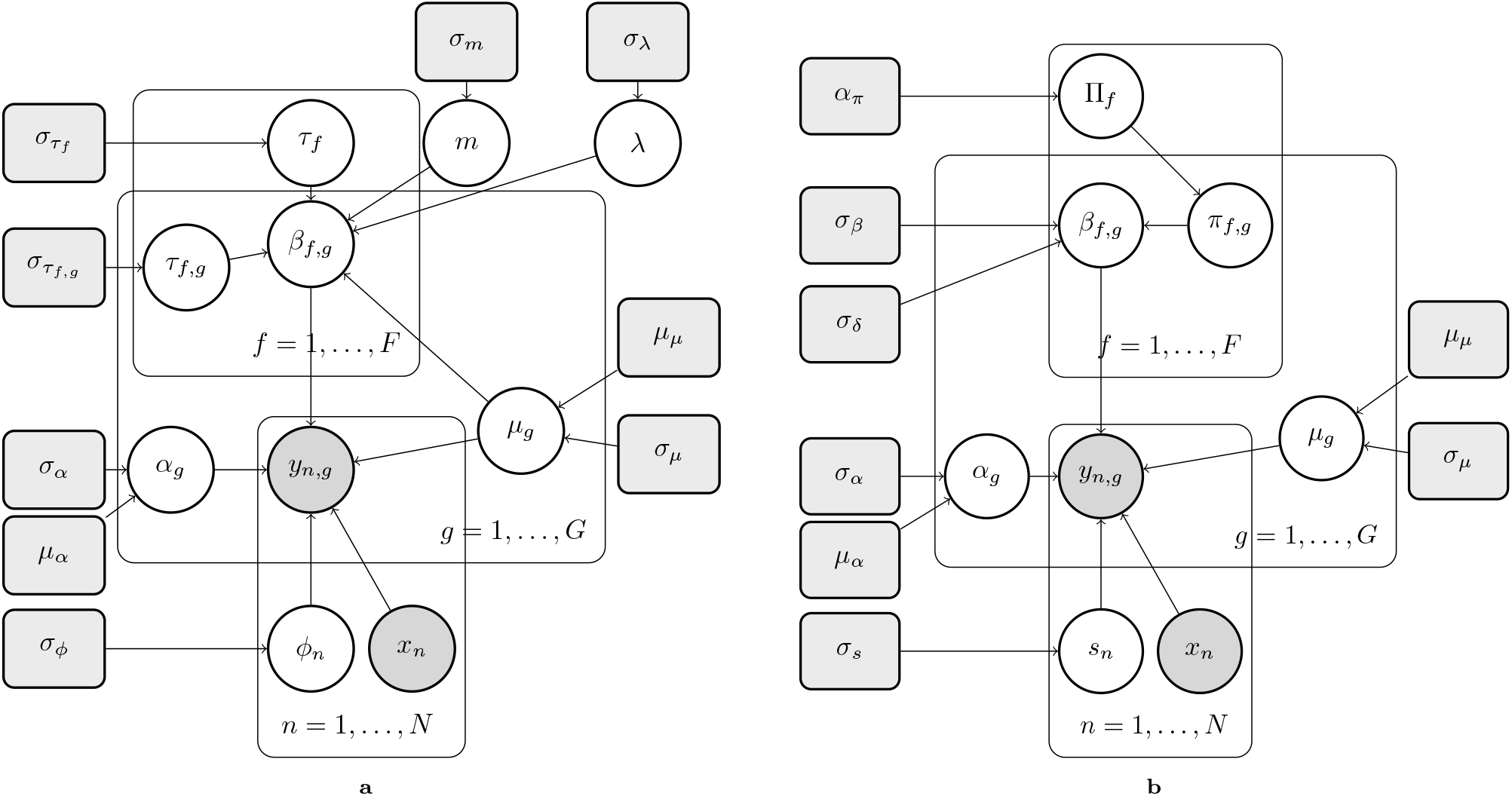
Probabilistic graphical models of the GLORB non size-factor (A) and size-factor (B) models. Hyperprior values are shown in grey boxes while parameters that are learned are put into circular nodes. Grey circular nodes are observed such as *y_n,g_* and *x_n_* which represent the observed counts and the sample specific information respectively. Plates represent repeating this structure across all iterations of a variable with the *n* plate representing each data point up to *N*, the gene plate representing each gene *g*up to *G*, and the *f* data plate representing each factor up to *F*.

**Table 1.**
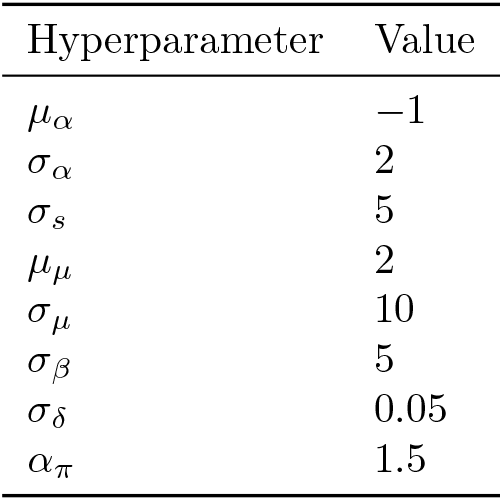
Hyperparameter values for the GLORB size-factor model.

| Hyperparameter | Value |
| --- | --- |
| $\mu_\alpha$ | -1 |
| $\sigma_\alpha$ | 2 |
| $\sigma_s$ | 5 |
| $\mu_\mu$ | 2 |
| $\sigma_\mu$ | 10 |
| $\sigma_\beta$ | 5 |
| $\sigma_\delta$ | 0.05 |
| $\alpha_\pi$ | 1.5 |

**Table 2.** Hyperparameter values for the GLORB non size-factor model.

| Hyperparameter | Value | Hyperparameter | Value |
| --- | --- | --- | --- |
| $\mu_\alpha$ | -1 | $\sigma_\alpha$ | 2 |
| $\sigma_\phi$ | 1 | $\mu_\mu$ | 2 |
| $\sigma_\mu$ | 10 | $\sigma_m$ | 1 |
| $\sigma_\lambda$ | 1 | $\sigma_{\tau_f}$ | 3 |
| $\sigma_{\tau_{f,g}}$ | 3 | $\epsilon$ | $\approx 0$ |

## 3 Results

### 3.1 Evidence of global shifts

We looked for evidence of global shifts from size factors due to changes in the median ratio from global upregulation as illustrated in Fig 2a in three datasets that have known global upregulation effects, as well as one dataset that we expected to have global upregulation effects. To identify the presence of global upregulation effects, we assume that the data-generating process has ensured that the technical contributions to total library sizes for samples are drawn from some common distribution across groups, i.e., if there is no global shifts present, the library size estimates from a method such as DESeq2 will be identically distributed for all groups; specifically, by the Central Limit Theorem we would expect the mean of library sizes estimated to be identical across groups in the limit of the sample size. However, if there is a global effect upon RNA transcription between groups, then the library size estimates for each group would be biased as the global upregulation is absorbed into these. If these assumptions hold, then performing a standard hypothesis test, such as the two sided Mann-Whitney U test, between the estimated group library size estimates will reveal the presence of global upregulation.

**Figure 2.**
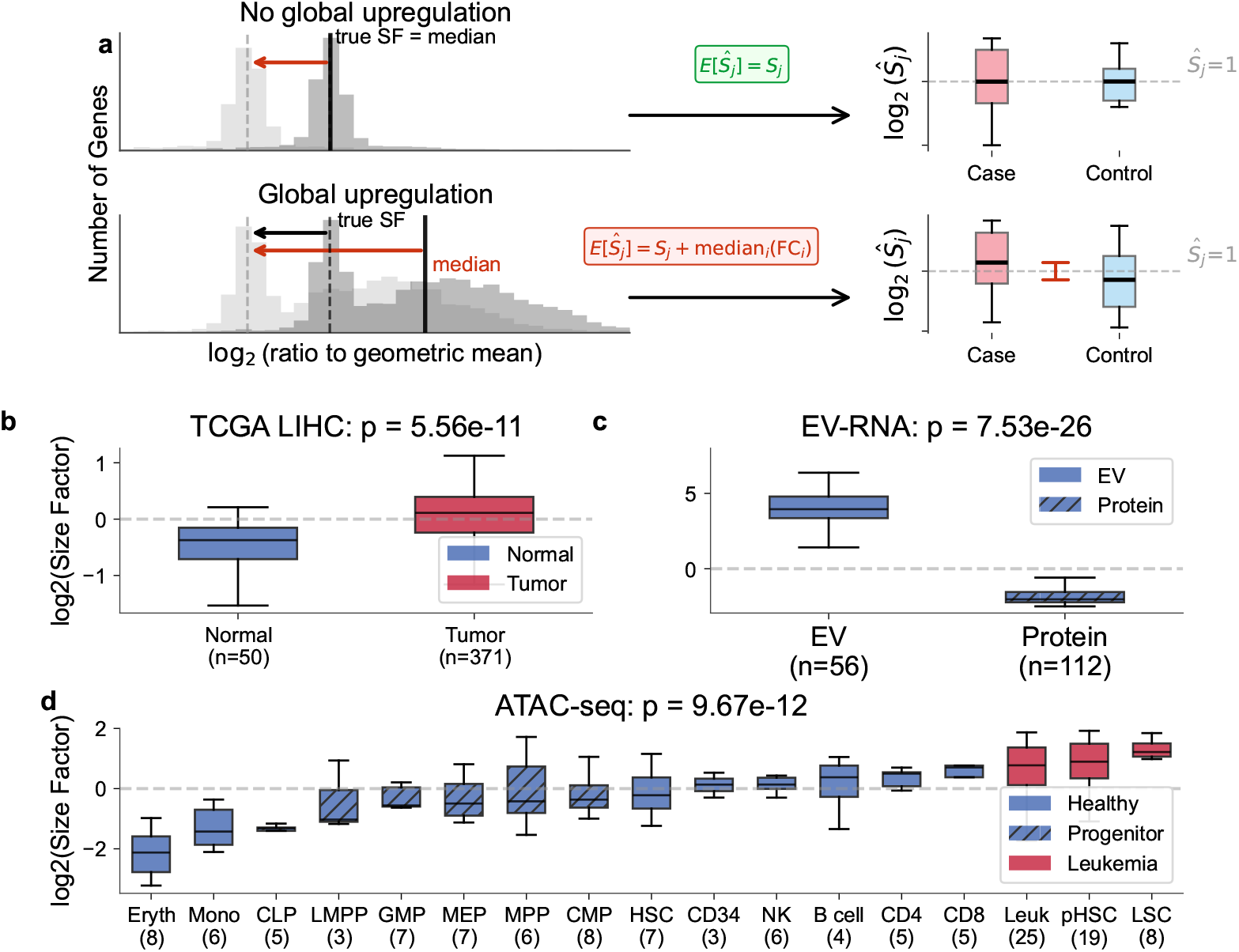
(a) Illustration of bias introduced by median-of-ratios normalization under global upregulation. Histogram of ratios of a genes’ counts in a single sample to the geometric mean of counts of that gene across samples. X axis showing ratio to the geometric mean of all samples for a specific gene. Dark line showing calculated median of ratios with observed data in dark histogram dashed line showing scaling to true scaled ratios in light gray. Red line showing scaling amount. Dashed dark line and black arrow showing true size factor and correct scaling amount. Top plot showing scaling to correct size factor effect in green showing boxplots with centered means with bottom plot showing the overscaling effect as labeled in red with means that are separated. (b) Dataset comparing the distribution of log size factors between LIHC Normal samples in blue and LIHC tumor samples in red. Significance tested with a Mann-Whitney U test. (c) Dataset comparing RNA-seq counts in putative Extracellular Vesicle fraction vs Protein fraction in a dataset of plasma previously collected. Box plots shown comparing distribution of log size factors calculated by DESeq2. Significant by ANOVA. (d) Dataset comparing ATAC-seq counts in peaks by cell type colored by healthy cell type, progenitor cell type and leukemia cell type. Number of samples in each group is labeled underneath the cell type label. Difference between mean log size factor significant by ANOVA. Box plots shown comparing distribution of log size factors calculated by DESeq2.

We acknowledge that these assumptions about the shared size factor distribution across groups may not always hold, or be estimable in small sample sizes, but we expect this to be true in cohorts such as the TCGA[44], where specific care was made to analyze the RNA seq counts uniformly, and where it has been previously shown that global upregulation exists [22, 46]. One such dataset with evidence for hypertranscription (an increase of transcripts when normalized by chromosome) provides evidence for the presence of global upregulation in Liver hepatocellular carcinoma (LIHC)[46]. We compare means of log(*s_n_*) between cancer and adjacent normals and find it significant at *p ≤* 0.001 (Figure 2b). We also compare across multiple different cancers in the Cancer Genome Atlas (Figure S5) in cancers with at least 25 healthy adjacent normals and find that 9 are significant (p*≤*0.01), with 4 being significant at (p*≤*0.001).

We examine another dataset that was processed with spike-in controls and analyzed RNA measurements in human plasma derived extracellular vesicles. The dataset contains multiple fractions processed by high pressure liquid chromatography of cell-free RNA. We combine fractions as shown in the paper to compare cell-free RNA from the Extracellular Vesicle fraction of plasma to the cell-free RNA collected from the protein fraction of plasma and find it is also significant when normalizing with the median of ratios method (p=7.53 *×* 10*^−^*^26^, Figure 2c).

For our final example dataset we analyzed cellular differentiation where there has been previously speculated to be global upregulation[32]. This dataset is also notably not an RNA-seq dataset but a dataset of reads in ATAC-seq peaks. Methods for RNA normalization are often used in ATAC-seq datasets as well and so we examine the difference in means of log(*s_n_*) between these two groups as well. We find that cell differentiation shows an expected effect where accessibility has an implied higher level in undifferentiated cell types due to its large estimated size factors (Figure 2d). In this dataset size factors may be masking global upregulation of transcripts in undifferentiated stem cells and global down regulation of transcripts in cell such as erythrocytes [6].

Given the potential presence of global upregulation, and observed bias in log(*s_n_*) across multiple different data types, and biological contexts we next moved to benchmark our model on a simulated dataset and then apply it to real datasets to see if we would be able to recover information that was obscured using current common normalization methods.

### 3.2 Model performance and application to real datasets

Next, we tested our model using a simple simulation framework described in Section 5.2. To examine our simulation result we first examined one sample from 3 different simulations and visualized the log_2_(counts+1) normalized data with PCA using centering and scaling in (Figure 3a–c) to ensure that sample types clustered and separated from each other. We see clear visual separation in the PCA space meaning we expect that we should see many differences in means between genes as originally intended.

**Figure 3.**
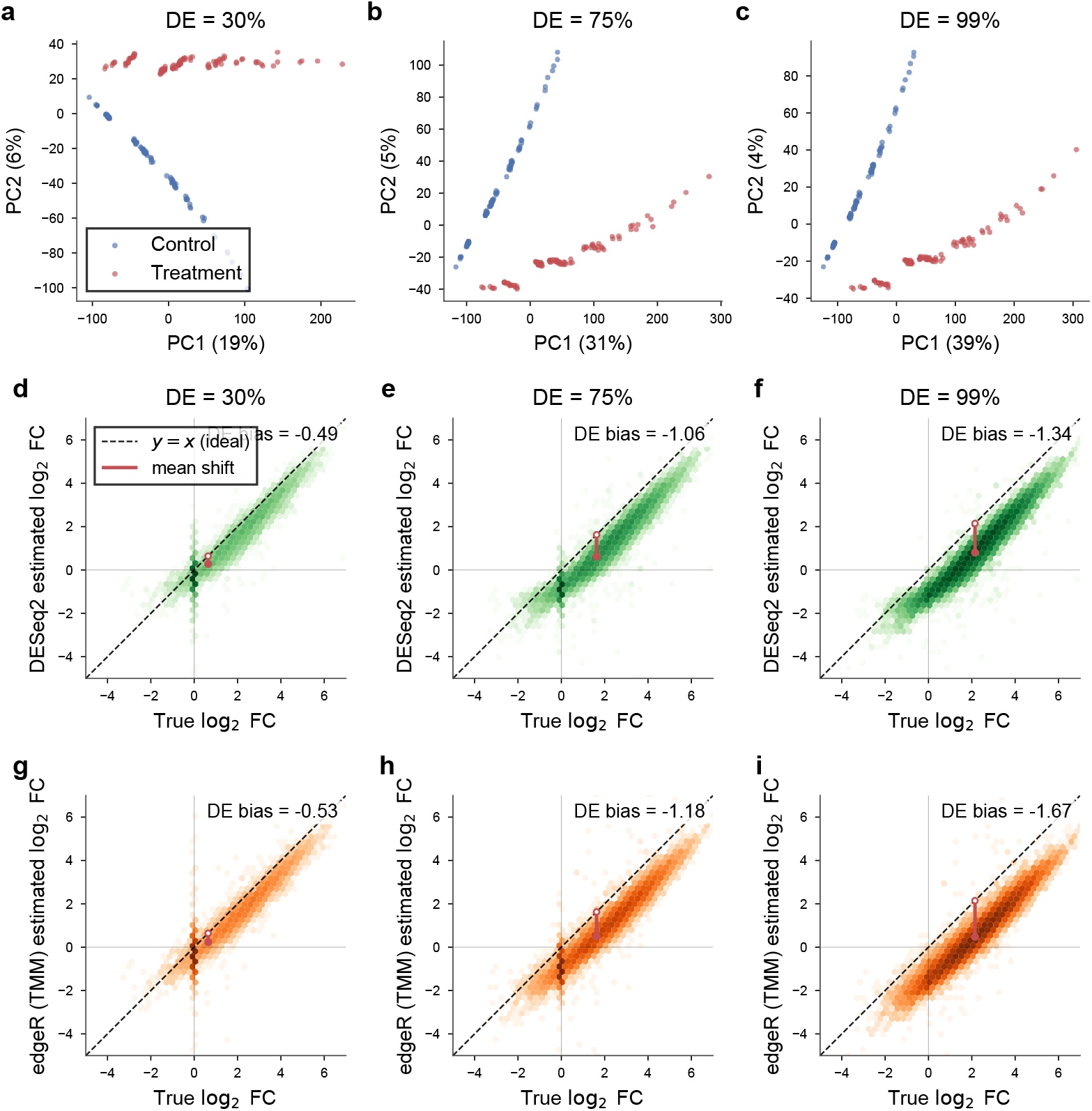
Benchmarking of GLORB. All panels use simulated datasets of 100 samples per group and 200 total (control and case) with the indicated fraction of genes defined as differentially expressed. The dark dot around the origin in the first two columns represents the genes that remained unchanged between conditions. Log of total number of points is plotted by color in each bin. A line showing the vertical distance between the means of both estimates and the y=x line where the estimates would be unbiased with the distance labelled as the DE bias. (a–c) PCA of simulated counts showing separation between control and case groups with 30% (a), 75% (b), and 99% (c) of genes differentially expressed. Counts are normalized as log_2_(counts+1); PCA is centered and scaled. (d–f) Density of DESeq2-estimated log_2_ fold changes versus true log_2_ fold changes with 30% (d), 75% (e), and 99% (f) of genes differentially expressed. (g–i) Density of edgeR-estimated log_2_ fold changes versus true log_2_ fold changes with 30% (g), 75% (h), and 99% (i) of genes differentially expressed.

Next we attempted to confirm that global upregulation will lead to bias in two current well known methods edgeR and DESeq2. We calculated the bias in predicted log fold change between case and control by comparing the true log_2_ fold change vs the DESeq2 calculated fold change and the edgeR calculated fold change. We observed a bias of −0.49, −1.06, and −1.34 for percentage of DE genes of 30% 75% and 99% respectively when using the DESeq2 normalization method (Figure 3d–f). We also observe a bias of the mean estimated log fold change from the true log fold change when using edgeR with biases of −0.53, −1.18, and −1.67 for percentage of DE genes of 30% 75% and 99% respectively (Figure 3g–i). This confirmed and illustrated our expectation that global upregulation is masked when using either median of ratios to normalize or a trimmed mean of M-values method to normalize out size factor differences.

We examined the performance of our model across our simulated scenarios with four other different methods designed to account for library size and look for differences in gene counts across genes. We also run our model on a dataset with ERCC spike-ins to calibrate it on a ground truth dataset, and run our model on cancer types that are known to have global upregulation but do not contain ERCC spike-in controls. In the simulation scenarios we pick two sample size numbers with complete simulation results for all simulated sample sizes shown in Figure S1 and Figure S2. For many simulations with no bias between samples but large global upregulation fractions and total sample numbers both the un-normalized t-test or mann-whitney have the highest median specificity of all methods (Figure S2(k,l)). Model fits for a single data simulation but with multiple different inference runs are shown in Figure S3 and Figure S4. We find that our model does not lose specificity as the percentage of genes that are differentially expressed increases Figure 4a. We also find that our model has the highest area under the receiver operating curve when comparing multiple across all cutoffs in different global upregulation scenarios (Figure S1(b–f), results calculated by one sided Wilcoxon signed rank test with *p ≤* 0.005).

**Figure 4.**
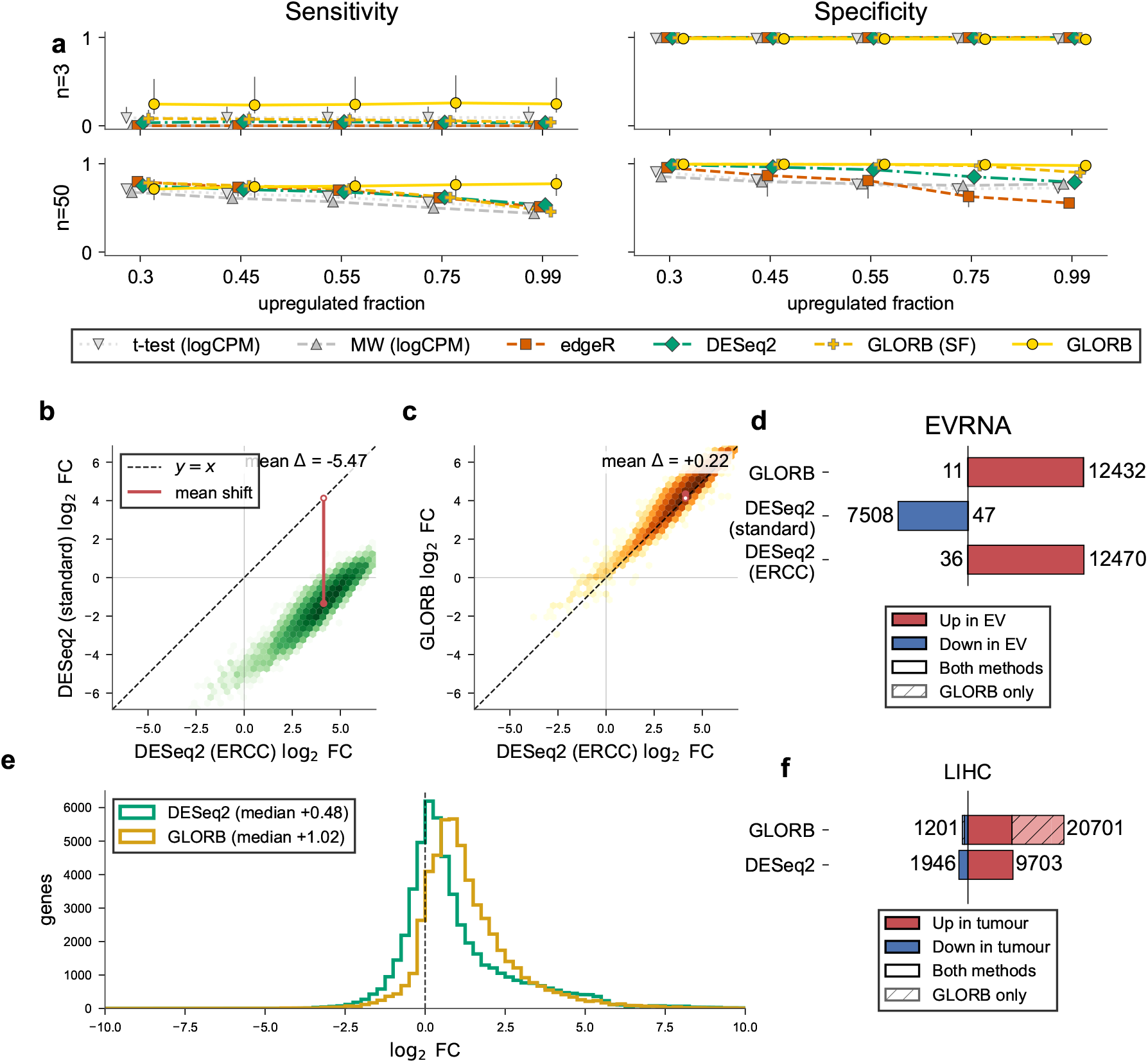
Benchmarking of GLORB. (a) Performance of both models of GLORB vs other normalization methods for library size across different sample sizes and percentage of variable (DE) genes. Sensitivity and specificity are calculated using the default method for both edgeR and DESeq2, with a 95% posterior likelihood of a gene being as high or higher than a fold change of 1 used for GLORB. t-test and Mann-Whitney performed on Log(counts*1e6+1) and using a p value of 0.05 to select after BH multiple testing correction. Black lines show minimum to maximum range across data generating seeds. (b) Density plot of estimated log_2_fold changes from DESeq2 of genes vs DESeq2 log fold change calculated with ERCC spike-in controls in a dataset comparing RNA from the putative Extracellular Vesicle fractions of plasma vs the putative RNA fractions of plasma with Δmean showing the difference between the estimated log fold change by each methods and the red vertical line illustrating the mean shift. (c) Density plot of estimated log_2_ fold changes from DESeq2 of ERCC spike-in controls vs the same estimated log fold changes using the no size factor GLORB model in a dataset comparing RNA from the putative Extracellular Vesicle fractions of plasma vs the putative RNA fractions of plasma with Δmean showing the difference between the estimated log fold change by each methods and the red vertical line illustrating the mean shift. (d) Horizontal barplot comparing genes found with GLORB non-size-factor model vs those found with DESeq2 both with and without using spike in controls for size factor normalization. For the GLORB barplot genes found by GLORB that intersected with those found by DESeq2 are plotted in dark colors while additional genes found by GLORB with changed means shown in light shaded colors. (e) Density histogram showing distribution of predicted log fold changes for the intersection of genes used by both methods in the LIHC dataset showing GLORB non size factor showing a majority as upregulated with an estimated modal log2 fold change that is higher than the same modal estimated log 2 fold change found using DESeq2. (f) Horizontal barplot comparing genes found with GLORB non-size-factor model vs those found with DESeq2 comparing LIHC and adjacent normal cancer samples from the Cancer Genome Atlas. For the GLORB barplot genes found by GLORB that intersected with those found by DESeq2 are plotted in dark colors while additional genes with changed means shown in light shaded colors.

We add an additional simulation comparison to test the effects of bias that are potentially introduced by sample processing and examine a potential failure mode of our model. In this failure mode we vary the average size factor between groups in our simulated data by examining size factors with an average that is 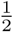 the size 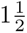 times larger and twice as large. We examine both specificity (Figure S7) as well as the shift of estimated log fold change (Figure S6). We show that our models retain comparable performance to models such as DESeq2 and edgeR across multiple upregulation fractions and total number of samples (Figure S7). Surprisingly despite being more susceptible to biases we find that our size factor model has the highest median specificity in 2 of the 3 scenarios for simulated datasets with at least 25 samples per group and a 0.55% or higher upregulation amount (Figure S7a).

Next, we examine our model’s performance in a dataset previously generated by our lab [18], in which there is a validated drastic shift in RNA content between conditions. In the experiment we analyze, RNA was extracted for sequencing from human plasma and fractionated into putative extracellular vesicle and non extracellular vesicle fractions. ERCC spike-in controls were added to all samples for normalization to control for technical variability. When comparing the ground truth, established via median of ratios normalization of ERCC spike-in control counts, with a median of ratios normalization performed with all genes, we find substantial bias with a difference of mean estimated log fold change of −5.47 between methods. This causes a majority of genes to be falsely estimated as down regulated. This is in contrast to our model, which yields results that closely match those obtained using ERCC based normalization Figure 4b–d.

Finally, we also compared our method to calculate differentially expressed genes in a variety of cancers (Figure S5). We focus especially on liver cancer in Figure 4 in particular because it has been previously shown to have hypertranscription [46]. In liver cancer we find our method detects a shift in the total upregulation of genes with the median gene having higher upregulation when measured by our method vs when estimated using DESeq2 (Figure 4). Where DESeq2 was found to have a significant difference between the distributions of the calculated size factors, we find a more than two fold increase in the total number of differentially expressed genes found (20,701 vs 9,703 calculated with DESeq2 Figure S5b) as expected. When reanalyzing datasets from TCGA we find many more upregulated genes while also finding a large overlap of genes that DESeq2 found as upregulated implying that global upregulation may be masked by size factors in analyses of these datasets.

## 4 Discussion

For the past 10 to 15 years, multiple papers have published results showing that global expression is an important and prevalent effect that exists across cell types, species, and conditions [22, 26, 31, 32, 42, 46]. Current best methods require adding spike-ins or are not readily available to be used with a full generalized linear model framework. Methods like ISnorm are specifically designed for cell specific size factors and require bulk size factors to already be calculated. We present what is to our knowledge one of the first methods to provide differential expression testing with full inference of a design matrix and linear model coefficients under global upregulation of RNA transcripts. We find signs that suggest global upregulation across TCGA [44] data, ATAC stem cell differentiation data, and see that this pattern is recapitulated in data sets with known global upregulation and spike-in controls. We expect our method will provide a window into analyzing previously collected RNA sequencing data sets as well as ATAC datasets, ChIP sequencing data sets, and others. We expect that with an easier to use, widely available computational method to detect and account for global upregulation, more complex biology can be investigated, including mechanisms such as drug response, cell differentiation, virus response, and developmental biology. We also expect that with further computational and hardware advancements it will be relatively easy to extend this methodology to single-cell sequencing data. Our methodology provides a Sensitivity Specificity potentially simple avenue to extend the way in which size factors are taken into account in RNA-seq analyses for both cells and bulk samples.

We recognize that our models and their implementation are limited in several ways. Particularly for the non-size factor model, we rely on the assumption that the size factors defining the libraries are drawn from a common distribution–that there is no bias due to technical preparation between condition and control samples. A second limitation is the reliance on the existence of marker genes. If there is massive upregulation and more than 99% of all genes are affected then as the number of genes that are marker genes goes to zero there will be an overdetermined system even with priors. This will make it functionally impossible to tell the difference between global upregulation between conditions and a true bias in PCR or other sample handling with our method. Finally, because this method uses variational inference to estimate distributions, the posterior uncertainty is underestimated. This means that it is very difficult to get accurate calibration for events that are very rare. If the goal is to filter out genes with an extremely low tolerance for false positives this method will be unable to determine between low likelihood and extremely low likelihood due to the light distributional tails on the normal distributions that it uses to create the mean field guide. However, seeing as this analysis scenario is unlikely to coincide with global upregulation, we do not believe it is a fundamental limitation.

This project could be easily extended to working with much larger datasets and there are a number of potential extensions that would allow it to scale in ways that are similar to scVI [24] while keeping some but not all of the statistical structure that it currently has. One notable way to increase throughput that is essentially identical to scVI would be to perform amortized inference similar to LD-VAE [40] or VEGA [36] project. Choosing the size of the latent space would be incredibly important however, in order to allow for the full linear model to still be learned. However, reducing the total dimensionality from 20,000 genes down to only 2,000, or 200 co-linear gene combinations would allow for a 10-100x increase in the total number of data points used. This would allow increasing datasets of 500 to around 50,000 on a laptop and increase datasets of around 5,000 to around 5 million data points. While this is still short of what is needed for quick analysis of massive datasets, it shifts towards the direction needed for modern single-cell analysis.

Another option for inference would be to perform parallel coordinate ascent variational inference with a modified version of the model that doesn’t use the negative binomial estimation for counts data and instead performs inference using the log normal distribution. Depending on performance losses that come from misreading variance off small gene counts this would allow extremely fast exact inference with modern hardware and would allow for much more optimization to occur.

## 5 Materials and Methods

### 5.1 Model inference and initialization

Models parameters are estimated using mean field variational inference through NumPyro [33] with the AutoNormal guide. Random seeds are handled with a NumPyro PRNGKey. Optimization is performed using ClippedAdam from the Optax library with a clipped global norm of 1.5, an initial learning rate of 0.01 and a scheduled exponential decay rate of 5 *×* 10*^−^*^4^ for 3,000 iterations. A full batch is used with no mini-batching for all experiments.

To allow SVI estimation of Bernoulli variables, we used a `RelaxedBernoulli` with a temperature of 0.3 in Numpyro [16, 27].

For initialization a novel mode-of-ratios initialization is used to find initial size factor estimations. The log ratio histogram is calculated where log(ratio) is between −7.5 and 7.5. 151 bins are calculated with edge bins corresponding to values with an absolute value greater than 7.5. To avoid artifacts from exact ratios due to sparse count genes only including counts of 1 we zero out the center bin centered at 0.0 and avoid it for future analysis. We then calculate the mode of the histogram across all bins by finding the bin with the largest value. This is calculated analyzing only bins that are not edge bins.

### 5.2 Simulation framework

To ensure model and coding specifications were met simulations were performed by constructing draws from a negative binomial distribution as implemented in the NegativeBinomial2 function in NumPyro where gene baseline means were drawn from the following distribution.

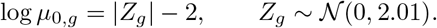

For *g* = 1*, …, G* genes. We then randomly picked a certain percentage of genes to be up-regulated genes in the condition samples. A gene *g* is designated as differentially expressed with probability *π*_DE_, i.e.,

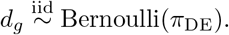

Then, for differentially expressed genes, a (natural) log-fold change is drawn as

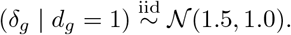

The condition-specific mean for gene *g* in sample *n* is then,

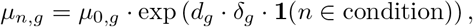

where **1**(*·*) is the indicator function and *n* = 1*, …, N*.

For our simulations, we calculate the dispersion formula in the same manner as DESeq2:

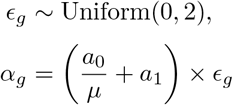

Where *µ* is the base mean, *ɛ_g_* is the gene effect, and *a*_1_ = 0.01 and *a*_0_ = 5. Then, size factors are generated for each sample:

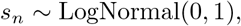

which then leaves the final simulation model for the observed counts of gene *g* in sample *n*:

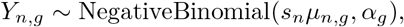

For our biased size factor experiments we draw the size factors from two separate distributions such that. For our control samples

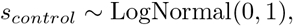

For our case samples the log normal distribution that size factors are drawn from is shifted so that

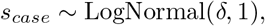

Where *δ* is defined as the amount of distribution bias that the case group has for all of its size factors, representing potential PCR bias or sample handling differences from the control group. We pick values such that *δ ∈* [*−*0.69, 0.41, 0.69] in the logarithmic scale. This is equivalent to a shift of 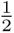, 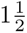 and 2.

For our simulation study, we use *G* = 20, 000 genes and consider different numbers of samples and upregulated genes;

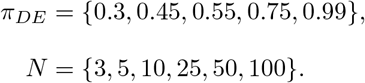

We use 10 random seeds for each scenario to assess the Monte Carlo variability between simulations, and infer each of these models with 5 separate seeds for each scenario. Comparisons are shown with the first seed of all inference runs.

### 5.3 Baseline methods, scoring, and evaluation

DESeq2 results for Figure 2 were calculated using pyDESeq2 [29] while full results in simulation were calculated for edgeR and DESeq2 using their R libraries with the most recent version. A BH adjusted p-value cutoff of 0.05 was used as the threshold for both DESeq2 and edgeR. edgeR estimates were calculated using the exact test. For datasets with sparse counts DESeq2 was run with the `poscounts` size factor estimation due to sparse data and a large sample number. Genes were counted as called as differentially expressed (DE) if the sign was concordant with the true Log Fold Change, as well as a log_2_ fold change of 1. For classical methods such as t-test and Mann-Whitney a false discovery rate correction was performed using `scipy.stats.false_discovery_control(’bh’)` [41]. T-tests and Mann-Whitney tests were calculated on log_2_(counts + 1) as a reasonable baseline to provide variance stabilizing. Log counts per million (CPM) versions were calculated as 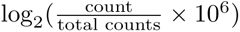. Mann Whitney tests are calculated two sided. For the GLORB method posterior likelihoods were calculated and a gene was only called as upregulated if it had at least a 95 percent chance of being as high as its estimated value or higher so that *p*(*|*LFC*| ≥* 1) *≥* 0.95. This ratio is calculated and is similar in style to ALDEx2 [11]. This was done to prevent extremely uncertain genes close to zero from counting as being differentially expressed.

### 5.4 Data sources and pre processing

All data sources used were publicly available and can be downloaded either from GEO or from Recount3. For Recount3 all cancer datasets from the TCGA were used with filtering to use sample types `01` for tumor or `11` for normal. Cancer datasets were filtered for those that had at least 25 normal samples. The EV-RNA dataset can be downloaded from GEO using accession GSE205301. ERCC genes were excluded from analysis except when rerunning and using them to calculate normalization coefficients. ATAC-seq hematopoietic cell datasets was downloaded from GSE74912.

### 5.5 Code availability

Anthropic models (Claude Opus and Fable) were used to assist with code writing and data analysis. Final versions of all scripts were audited by the authors to ensure correctness and then run to generate the results and figures. To facilitate an efficient code audit of the performance experiments, a centralized performance calculation script was used `calculate_scores.py` with all except two plotting scripts only using outputs from this script.

Publicly available data was downloaded from the Gene Expression Omnibus or GEO. For TCGA data, counts were downloaded using the Recount3 package [45] and metadata was also downloaded using this package. Scripts for the preprocessing of all data and graphing of the figures used in the manuscript is available at https://github.com/rowancallahan/global_upreg_seq_manuscript. Both models can be found and used from the repository available at https://github.com/rowancallahan/global_upreg_seq

## Acknowledgments

The authors would like to thank Ruby Fore for her insightful discussion of the Central Limit Theorem in its relation to this work. The authors thank Prof. Jordan G. Bryan for detailed comments on the clarity and presentation of the model formulation. The authors thank Prof. Galip Gurkan Yardimci, Prof. Olga Nikolova, and Dr. Travis Moore for comments and discussion. Finally, the authors thank Shuhao Wei for his feedback on model initialization methods. Anthropic models (Opus and Fable) were used to provide line edit and grammar suggestions.

## Author contributions

R.C. conceived the project, developed and implemented the models in code, conducted all simulation studies and real-data analyzes, selected and curated public datasets for benchmarking, produced all figures, and wrote the initial manuscript draft. S.D.C. contributed to conceptualization and formalization of the statistical model, including the probabilistic framework, and prior specifications; provided guidance on simulation study design and interpretation; reviewed and approved the selection of public datasets for benchmarking; provided feedback on the manuscript including structural reorganization, literature recommendations, and methodological framing; and contributed to conceptualization throughout the project. Prof. T.T.M.N. provided mentorship and scientific guidance, edited the manuscript, suggested the idea of a model including size factors and suggested the inclusion of ERCC spike-in data as a validation benchmark.

## Supplementary figures

**Figure S1.**
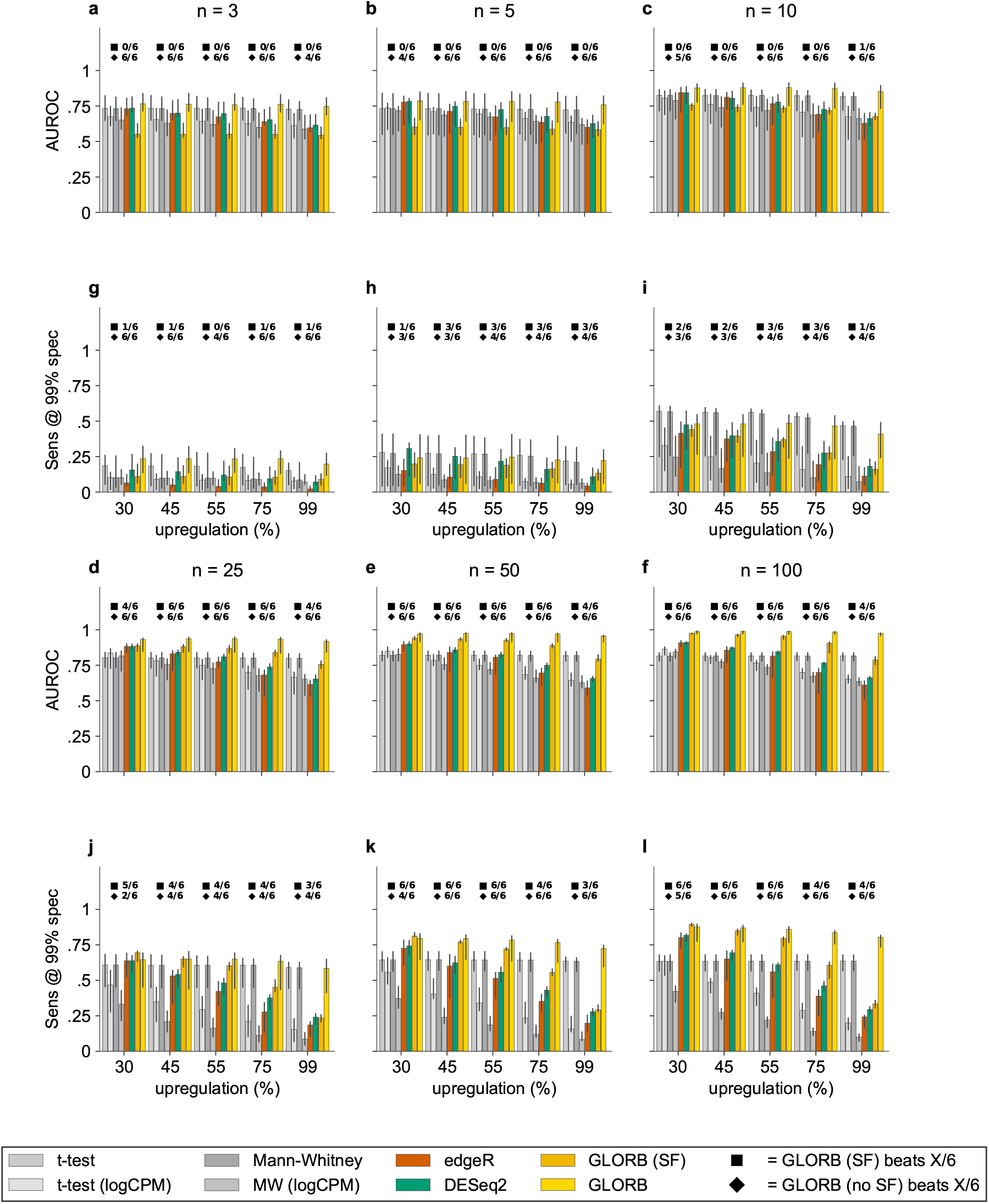
Comparison of performance of both AUROC and sensitivity at 99% specificity. Each bayesian model shows only data from the first inference run for each permutation. (a–f) Barplots with ranges showing median performance for a method and range from highest to lowest performance for each resampling of data. Barplots show comparison of number of non bayesian models that GLORB-SF and non size factor model have a higher area under the receiver operating curve out of all 6 comparisons with each of the other models used. Each plot starts at 30% differentially expressed genes and increasing to 99% differentially expressed genes. Columns show increasing number of samples used per comparison group. Non size factor model shown with a diamond and size factor model shown with a square. (g–l) Same comparison strategy as used in (a–f) but instead comparing sensitivity at estimated 99% specificity by interpolating along the receiver operator characteristic for each of the models. Non size factor model shown with a diamond and size factor model shown with a square.

**Figure S2.**
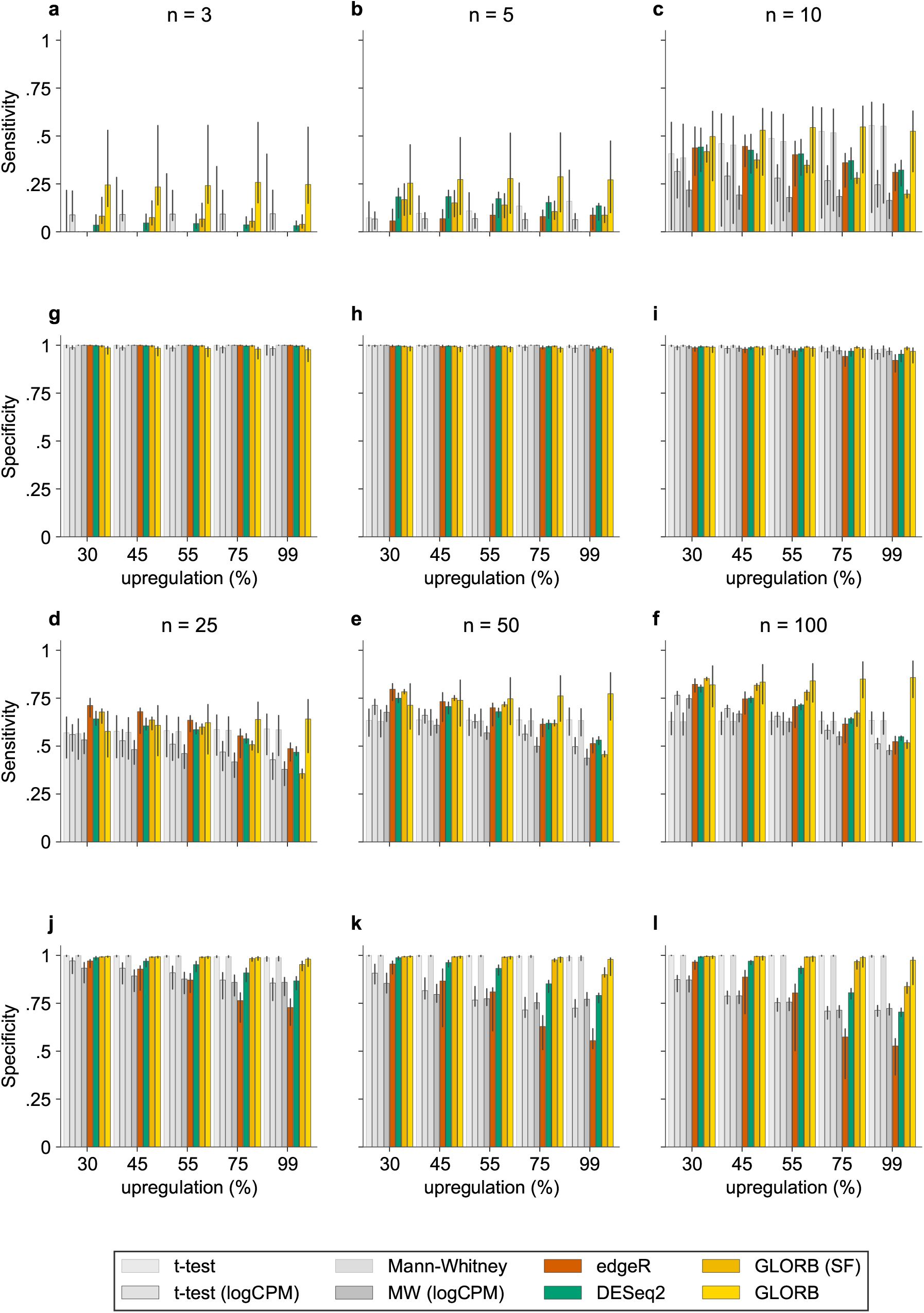
Barplots with ranges showing median performance for a method and range from highest to lowest performance for each resampling of data. Extended sensitivity (a–f) and specificity (g–l) performances calculated against simulated data for ground truth with called significant for classical methods using *P ≤* 0.05 after BH correction and estimated log fold change *≥* 1. For our Bayesian model we picked genes with a 95% posterior interval being higher or equal to our cutoff in our model. The number of samples in each condition ranged from 3 per group to 100 per group. Percentage of differentially expressed genes ranged from 30% (a,f) to 99% (e,j). Each bayesian model shows only data from the first inference run for each permutation.

**Figure S3.**
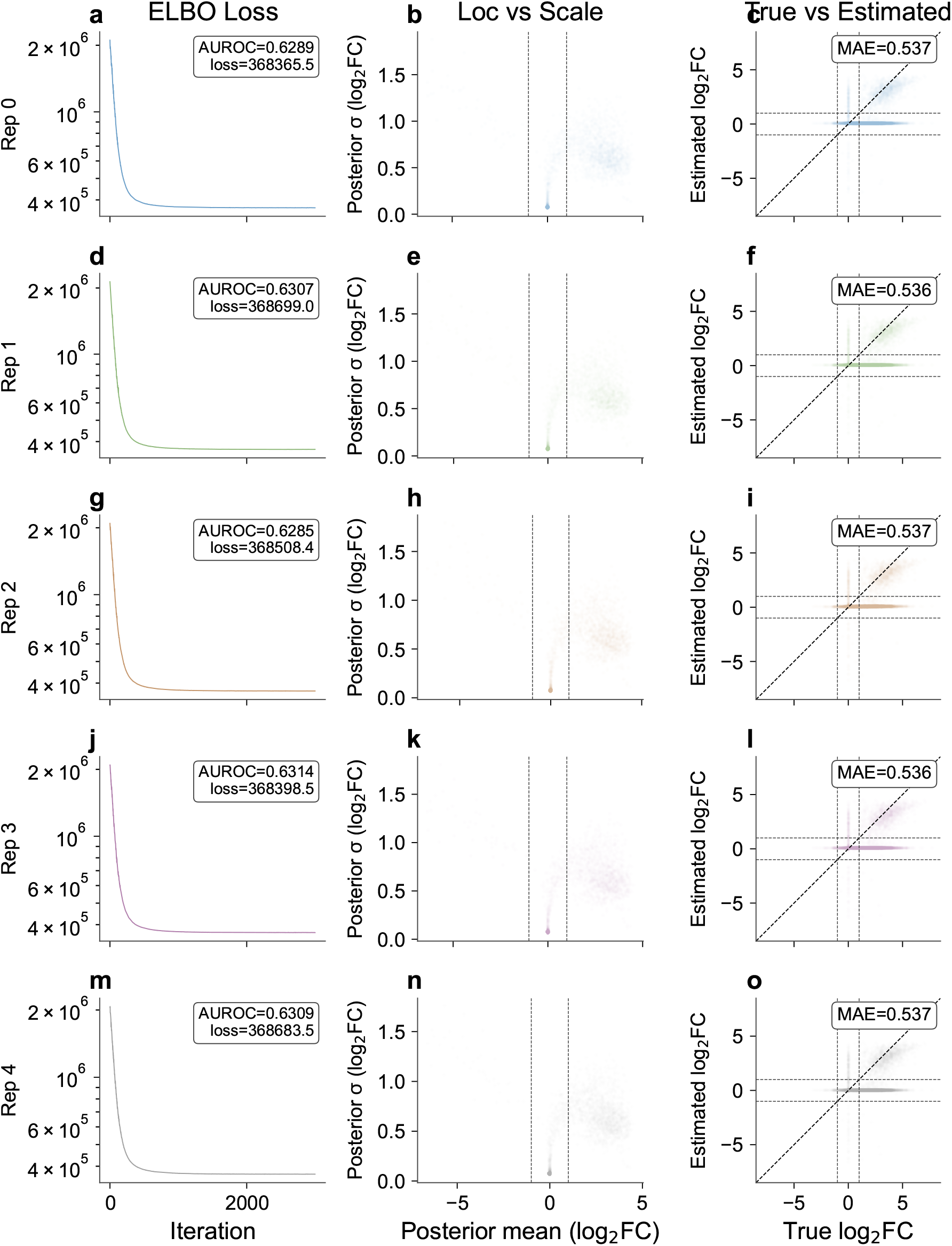
Comparisons across multiple iterations of optimization of the size-factor model of the same scenario showing consistent behavior (a,d,g,j,m) of loss and final AUROC after 3000 iterations of optimization. (b,e,h,k,n) Plots showing posterior estimate of mean (Loc) and variance (Scale) for guides estimated with SVI. (c,f,i,l,o) Mean absolute error between predicted and estimated fold changes across optimization iterations of the same scenario.

**Figure S4.**
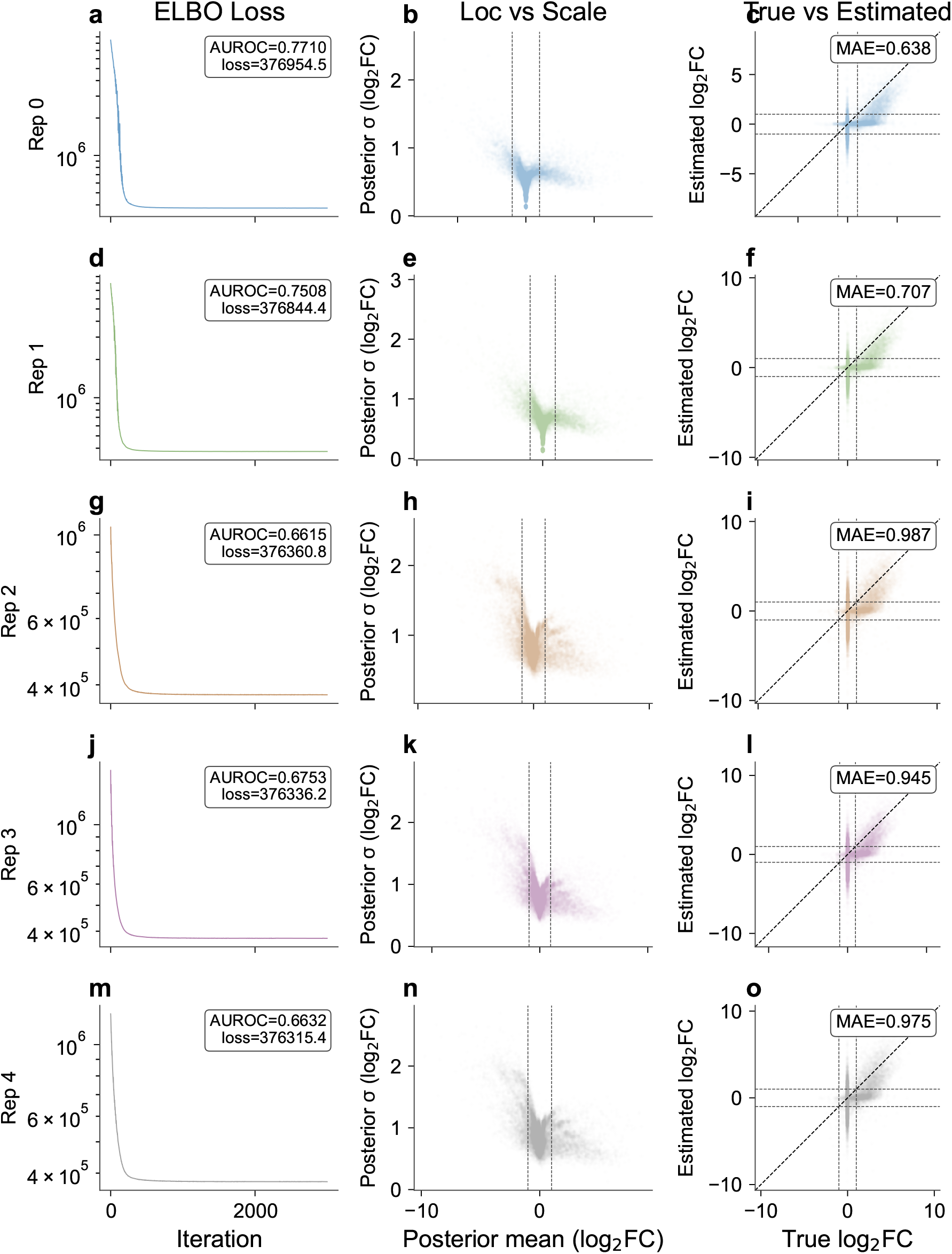
Comparisons across multiple iterations of optimization of the non-size-factor model of the same scenario showing consistent behavior (a,d,g,j,m) of loss and final AUROC after 3000 iterations of optimization. AUROC shows lower performance on three iterations than others with poorer convergence. (b,e,h,k,n) Plots showing posterior estimate of mean (Loc) and variance (Scale) for guides estimated with SVI. (c,f,i,l,o) Mean absolute error between predicted and estimated fold changes across optimization iterations of the same scenario.

**Figure S5.**
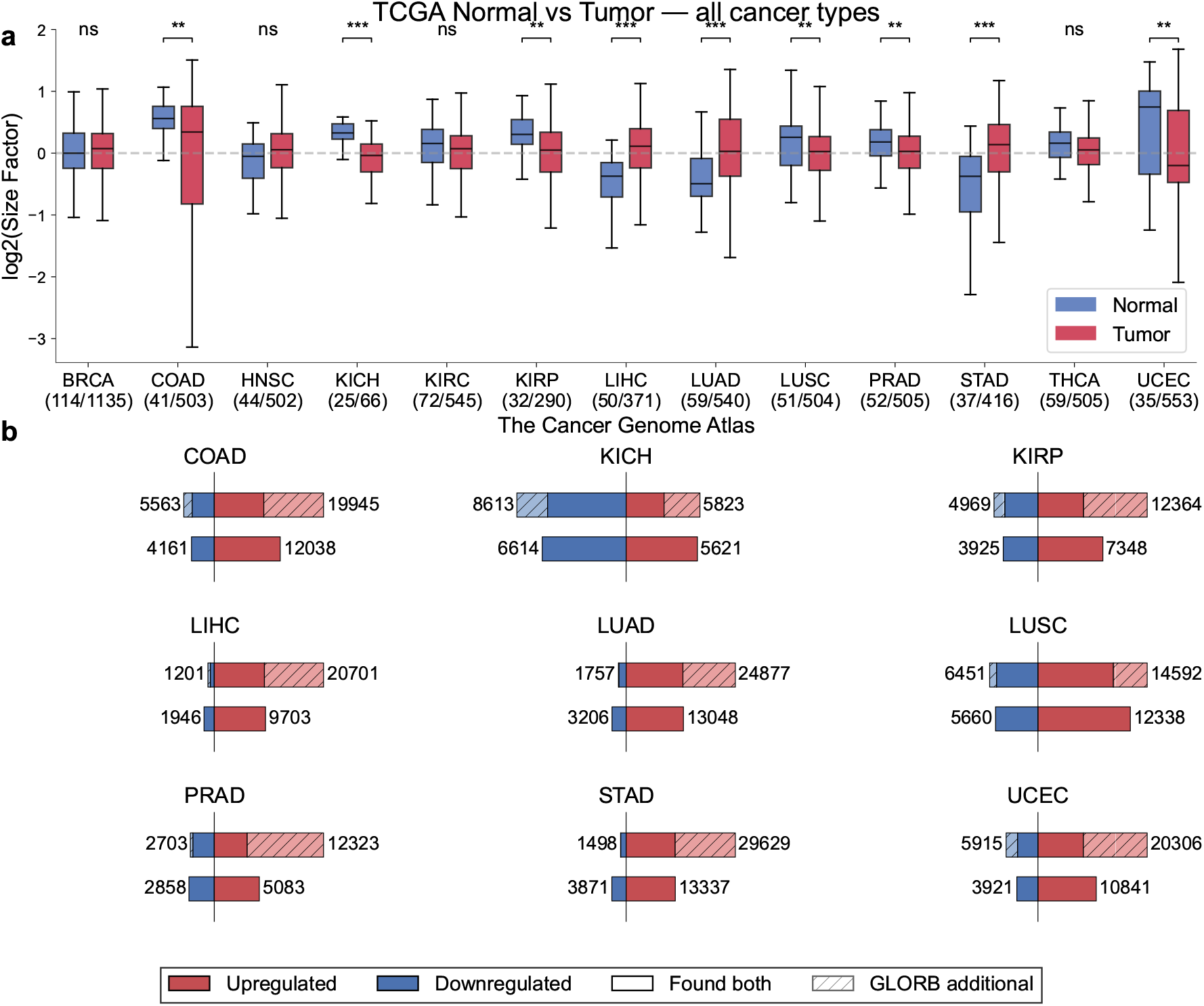
(a) Boxplots showing the comparison between log normalized pyDESeq2 estimated size factors for both tumor (red) and adjacent normal tissue (blue) showing only cancer types with at least 25 adjacent normal. Significance ** *p ≤* 0.01 and *** *p ≤* 0.001 all levels above called as N.S.(not significant). (b) Barplots comparing significant genes called with DESeq2 vs our non size factor model with the bottom plot showing gene counts increased (red) and decreased (blue) between conditions found in DESeq2, and the top plot showing the same increase and decrease. Dark colored portions of the bar in the top plot are intersections where genes found by our non size factor GLORB model are the same as those found by DESeq2, light shaded portions are those that are found only by our model.

**Figure S6.**
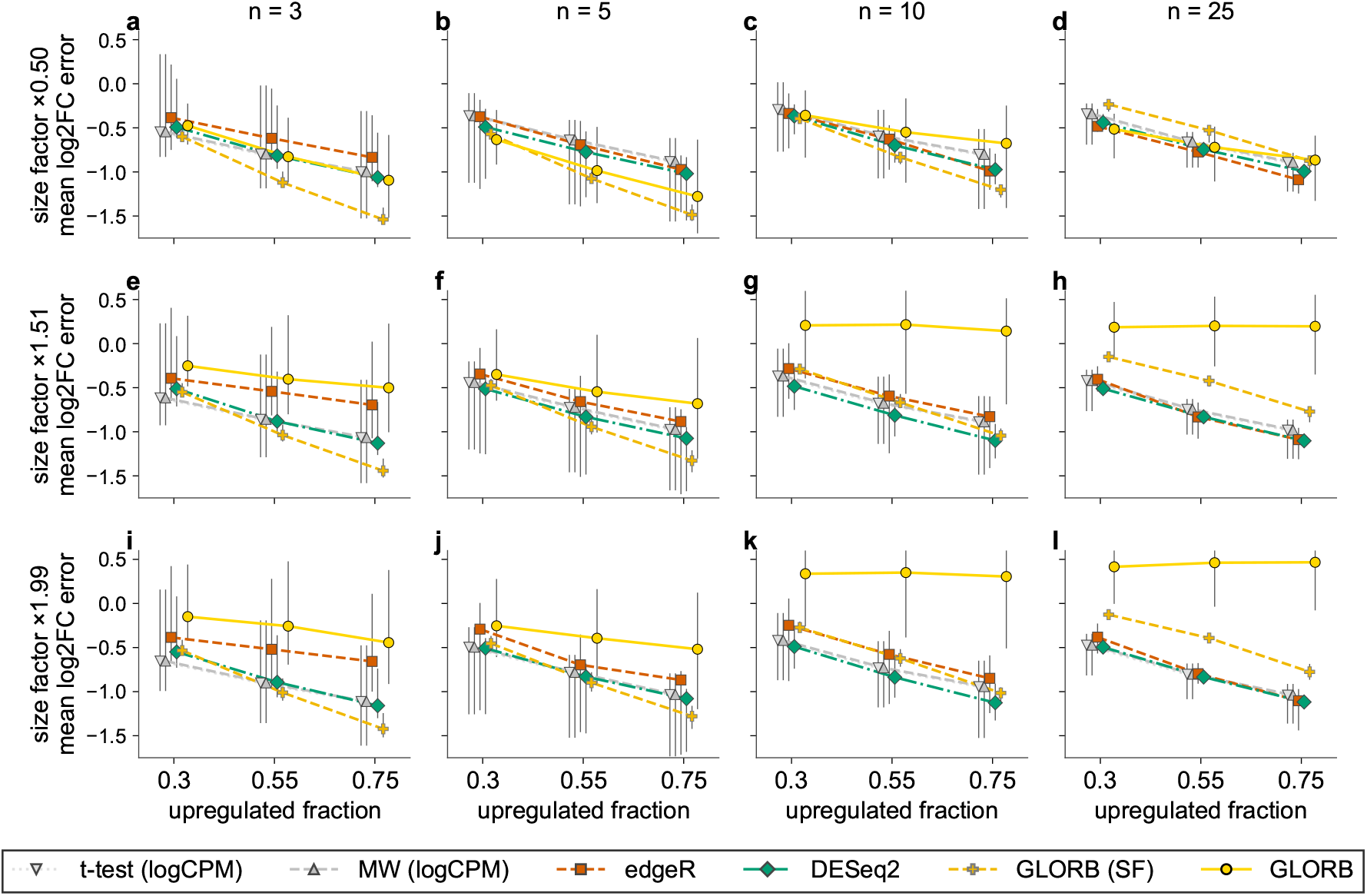
(a–l) Lines showing comparison of our 6 comparison models for differential expression at three separate bias levels for size factors where the distribution that each size factor was drawn from was biased relative to the control condition. Dots show median across all simulations of mean *log*_2_ fold change error. Line ranges show the range of the estimated mean difference from each calculated difference given a specific simulation from data simulator in a specific scenario. Performance of Bayesian models show the first inference of the total of 5 inference runs performed for each resampling of the data in a given simulation.

**Figure S7.**
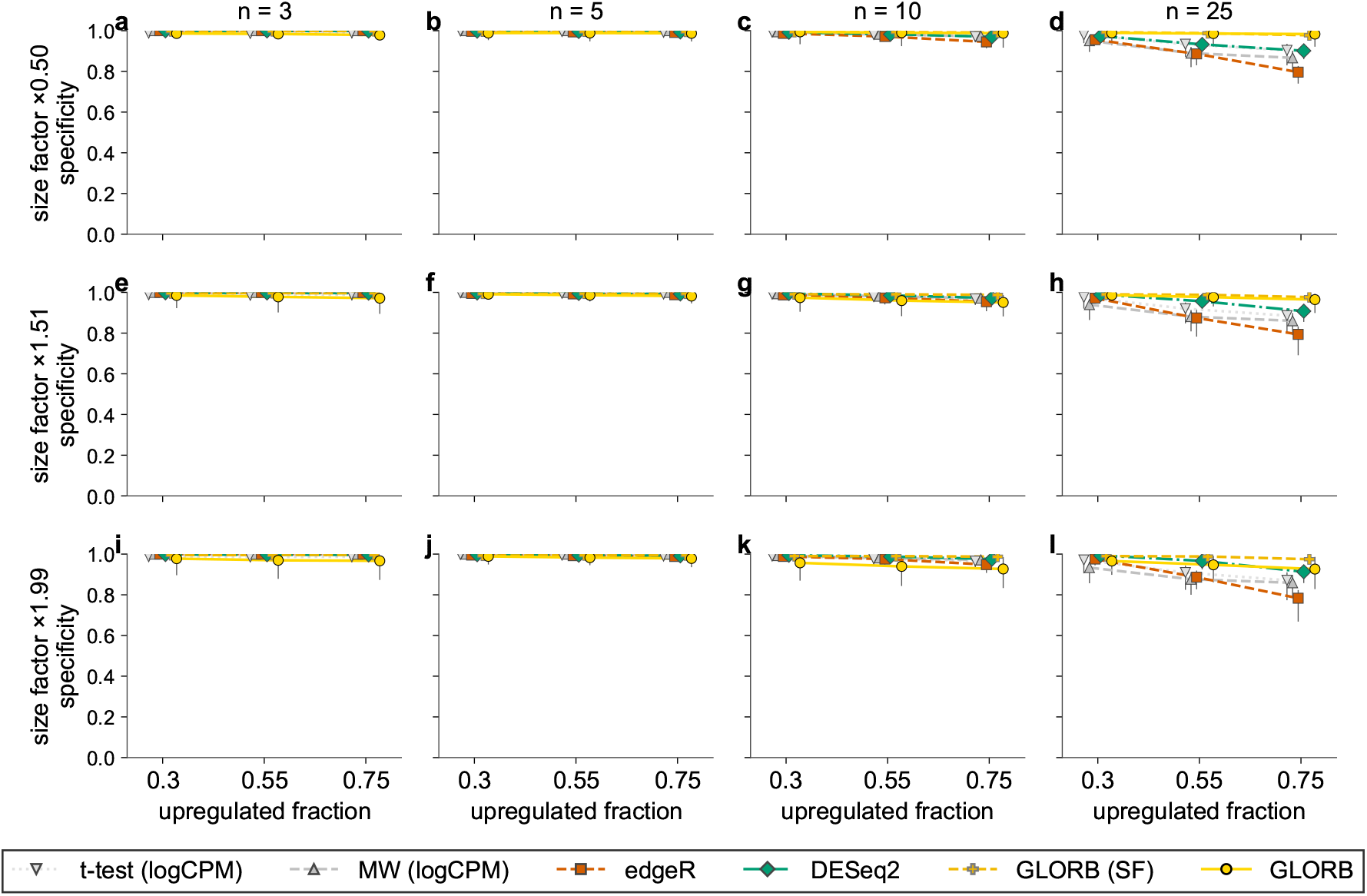
Lines showing performance of 6 models including both the size factor and non size factor model at multiple different fractions of upregulated genes as well as total number of samples per group. Performance is shown similar to in Figure S6 with ranges showing range across all different seeds for data creation but only using the first numpyro model inference seed for each model.

